# Stable transformation of *Micractinium conductrix* SAG 241.80: New tools for exploring photosymbiotic interactions

**DOI:** 10.64898/2026.01.27.702003

**Authors:** Fiona R. Savory, Victoria Attah, Estelle S. Kilias, Thomas A. Richards

## Abstract

Endosymbiosis played a key role in the evolution of cellular complexity, but mechanisms underpinning establishment and maintenance remain unclear. *Paramecium bursaria* harbours Chlorellaceae algal endosymbionts and is a valuable model system for understanding photosymbiosis. However, there are a lack of molecular tools available for exploring the association. Here we report stable transformation of the *P. bursaria* endosymbiont *Micractinium conductrix* SAG 241.80. Bioluminescent reporter assays were used to identify endogenous regulatory sequences for transgene expression and to refine a protocol for delivery of exogenous DNA. The bleomycin resistance gene *Shble* was then identified as an effective selectable marker for isolating transformed cell lines. We demonstrate that endogenous introns enhance transgene expression and that isolation of transformants with desirable characteristics can be achieved without extensive screening by using a 2A peptide to couple expression of an upstream non-selectable transgene to expression of the *Shble* selectable marker. Finally, we demonstrate that *M. conductrix SAG 241*.*80* transformants expressing fluorescent proteins can be introduced into host cells and observed in the absence of selection, enabling distinct endosymbiont genotypes to be unambiguously identified, compared and monitored within the host environment. The development of molecular tools reported here opens new avenues for addressing unresolved questions in photosymbiosis.

**Significance Statement:**

- *Paramecium bursaria* is a single-celled, mixotrophic ciliate which forms photosymbiotic interactions with Chlorellaceae green algae and is a popular model system for symbiosis research.
- We established a stable transformation protocol for the *P. bursaria* algal endosymbiont *M. conductrix* SAG 241.80 and identified properties of synthetic gene constructs which facilitate isolation of transformants with robust transgene expression that can be detected within the host environment.
- This marks significant progress in model system development and opens new avenues for exploring molecular and cellular mechanisms underpinning photosymbiotic interactions.

## Introduction

Endosymbiosis played a key role in the emergence of eukaryotic cellular complexity and is an important driver of evolutionary innovation (Archibald, 2015). Model systems for endosymbiosis in which host and endosymbiont cells form stable interactions but can also survive independently can provide valuable insights into intermediate stages during the transition from facultative associations to integrated organelles (Husnik and Keeling, 2019; Boudreau *et al*., 2025). Exploiting the full potential of such models to explore the molecular and cellular mechanisms underpinning endosymbiosis requires both partners to be amenable to genetic manipulation.

The freshwater ciliate *Paramecium bursaria* forms photosymbiotic interactions with green algae predominantly belonging to the Chlorellaceae family (Trebouxiophyceae, Chlorophyta). The symbiosis is based upon nutrient trade, exchange of carbon dioxide for oxygen and protection from natural enemies (Brown and Nielsen, 1974; Kawakami and Kawakami, 1978; Reisser, 1980; Albers *et al*., 1982; Sørensen *et al*., 2020). *P. bursaria* cells can contain several hundred algae which are each individually compartmentalised within a symbiosome, also called the perialgal vacuole. *P. bursaria* is a popular model system for investigating photosymbiosis largely due to the ability to separate host and endosymbiont cells and to re-establish the endosymbiotic interaction. Genomic and transcriptomic data are available for host and endosymbiont strains (Blanc *et al*., 2010; Kodama *et al*., 2014; Arriola *et al*., 2018; Leonard *et al*., 2025a, 2025b) and expression of *P. bursaria* genes can be altered via RNA interference (RNAi), allowing host mechanisms involved in establishing, maintaining and policing the interaction to be explored (He *et al*., 2019; Jenkins *et al*., 2021a, 2021b). However, the potential of the *P. bursaria* system for investigating molecular and cellular processes underpinning photosymbiotic interactions is hindered by a lack of genetic tools available for the algal endosymbionts. Furthermore, studying the fate of individual endosymbionts is limited because there are currently no methods available for tracking sub-populations of the endosymbiotic algae within host cells. Developing approaches to achieve stable heterologous expression in the algae would create new opportunities for probing the interaction, for instance through labelling endosymbionts with fluorescent reporters to monitor colonisation dynamics and the fate of distinct genotypes under different environments or host RNAi treatments.

Transformation and targeted genome editing have been reported for several Chlorellaceae algae (Run *et al*., 2016; Kumar *et al*., 2018; Lin and Ng, 2020; Kim *et al*., 2021; Gu *et al*., 2023; Kasai *et al*., 2024; Cui *et al*., 2025). However, studies are often limited by low efficiency and reproducibility, and species- and/or strain-specific challenges can be encountered, for instance due to compositional variation in the rigid and complex carbohydrate-rich cell walls which create effective barriers against delivery of exogenous molecules (Yang *et al*., 2016; Muñoz *et al*., 2018; Ortiz-Matamoros *et al*., 2018). In this study, we established a stable transformation protocol for *Micractinium conductrix* SAG 241.80, a Chlorellaceae alga originally isolated from *P. bursaria*. We then demonstrated the utility of this transformation approach by verifying that transgene expression can be detected in an endosymbiotic context within host cells.

## Results

### Endogenous regulatory sequences drive nuclear transgene expression

We used NanoLuc^TM^ luciferase (Hall *et al*., 2012) to develop and refine a transformation protocol for *M. conductrix* SAG 241.80. This highly sensitive bioluminescent reporter can be detected at very low levels and enables rapid screening of population-wide transgene expression based on analysis of cell lysates (Booth *et al*., 2018). Four vectors were designed and synthesised which each contained a codon-optimised *nanoluc* gene fused between endogenous *M. conductrix* SAG 241.80 noncoding sequences that flank nuclear-encoded genes (*ef1α, lhcII, psaD* and *rbcS*; Supplementary Table 1). As promoters and terminators have not been mapped for *M. conductrix* SAG 241.80, we used flanking sequences of 1 kb in length (including UTRs), reasoning that regulatory elements that could be used to drive transgene expression were likely to be present within these regions. After numerous attempts at transforming *M. conductrix* SAG 241.80 with different vector preparations and delivery conditions, low levels of luminescence (an overall indicator of transformation success) were detected in the cell lysates of populations electroporated with a single 3.5 msec, 6.0 kV/cm square-wave pulse and 40 µg of linearised vector. Although effect sizes were variable, *nanoluc* expression was detectable within the cell lysates of populations electroporated as above with all four vectors (Figure 1A), demonstrating that the tested combinations of endogenous noncoding sequences can all drive transgene expression in *M. conductrix* SAG 241.80.

**Figure 1.**
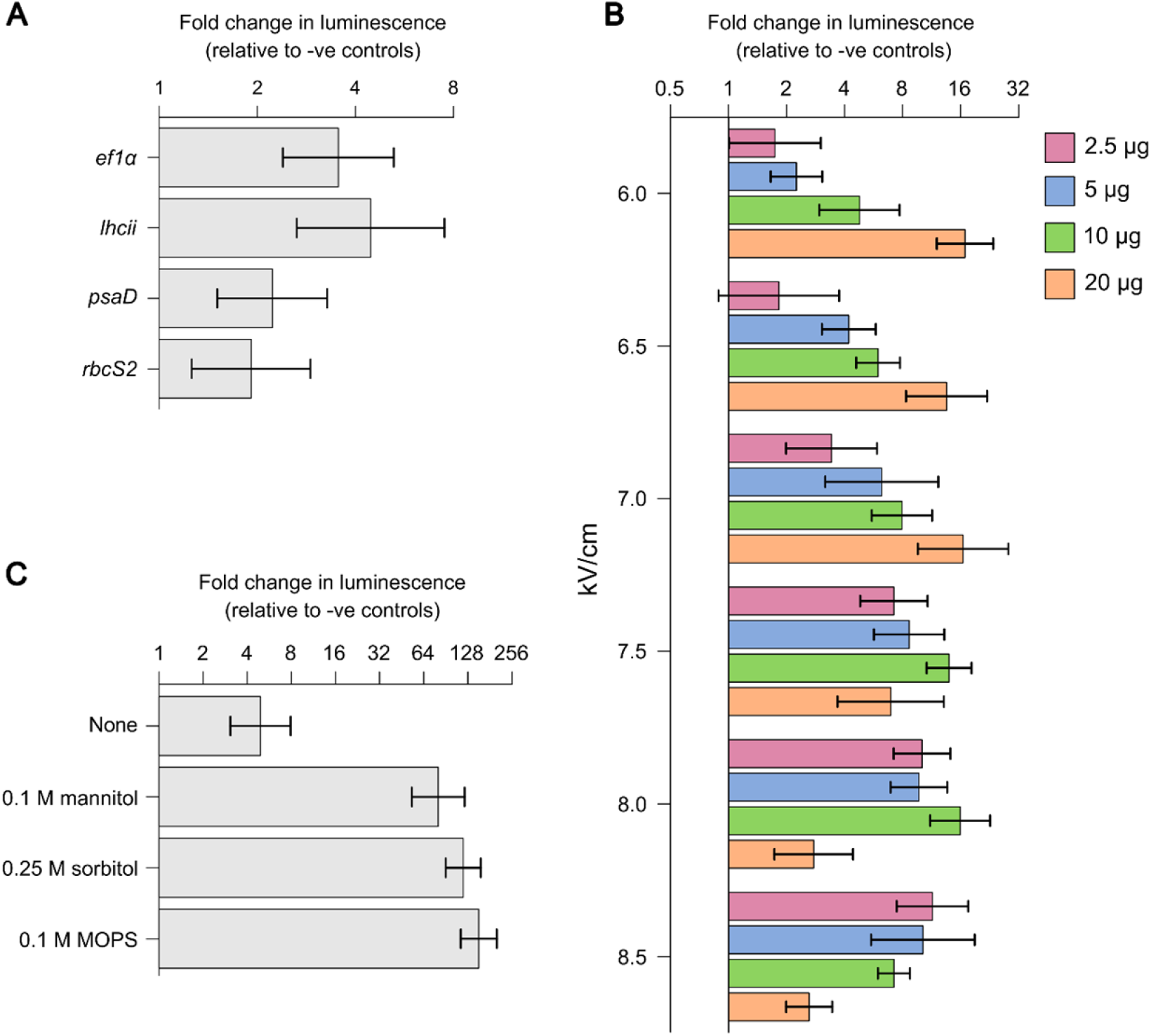
Development and refinement of a transformation protocol based on population level expression of NanoLuc^TM^ luciferase. (A) Vectors containing endogenous noncoding sequences which flank the coding regions of *M. conductrix* SAG 241.80 genes (*ef1α, lhcII, psaD* and *rbcS*) drive expression of a codon-optimised *nanoluc* gene. (B) Effects of electroporation field strength (kV/cm) on population level expression of NanoLuc^TM^ luciferase (an overall indicator of transformation success) are dependent upon the concentration of vector DNA supplied (coloured – see key). (C) Preculturing of *M. conductrix* SAG 241.80 populations in hyperosmotic growth conditions for 48 hrs prior to electroporation increases population level expression of NanoLuc^TM^ luciferase. Data are presented as fold change relative to controls electroporated without vector DNA and error bars represent 95% confidence intervals. Effects were considered as significantly different from controls when confidence interval error bars did not overlap with 1.

We sought to refine the transformation protocol through successive modification of the initial conditions described above. We used the *nanoluc* vector containing *lhcII* flanking sequences since this led to the highest mean luminescence signal (Figure 1A) and because *lhcII* was identified as the most highly expressed *M. conductrix* SAG 241.80 gene in a publicly available transcriptome dataset (Arriola *et al*., 2018; NCBI accession number GSE98781). Luminescence was detected in the cell lysates of populations electroporated using a single 5.0 msec square-wave pulse with field strengths ranging from 6.0 kV/cm to 8.5 kV/cm (Figure 1B). High quantities of linearised vector were required for robust luminescence signals to be detected at the lowest field strengths tested, indicating inefficient delivery of exogenous DNA under these conditions. Notably, robust luminescence signals were achieved with a moderately low quantity of linearised vector (2.5 µg per transformation) when high field strengths (e.g. 7.5 - 8.5 kV/cm) were applied, demonstrating that these electroporation parameters enabled effective delivery of exogenous DNA into cells and suggesting this process does not excessively compromise viability. Using 2.5 µg of linearised vector and a 5.0 msec, 8.5 kV/cm pulse as a baseline, small improvements in population level *nanoluc* expression were observed after supplementing the electroporation buffer with 40 mM sucrose (Supplementary Figure 1A) and reducing the volume of cell suspension used during electroporation from 100 µl to 50 µl whilst keeping total cell numbers constant (Supplementary Figure 1B). More striking increases in luminescence were detected when cells were precultured in hyperosmotic growth conditions for 48 hrs prior to electroporation (Figure 1C). These results reflect observations in other Trebouxiophyceae algae, as growth in media containing 0.6 M mannitol or 0.6 M sorbitol was a prerequisite for successful transformation of two *Coccomyxa* strains (Tatara *et al*., 2020). All subsequent *M. conductrix* SAG 241.80 transformations were performed using cells which had been precultured for 48 hrs in media containing 0.1 M MOPS (3-(*N*-morpholino)propanesulfonic acid), which led to an approximately 150-fold increase in mean luminescence values relative to background levels (Figure 1C) and was the least inhibitory to population growth of the hyperosmotic conditions tested (Supplementary Figure 2).

### Introns enhance nuclear transgene expression

We next set out to identify a suitable selectable marker for progressing genetic tool development. We examined sensitivity of *M. conductrix* SAG 241.80 to three antibiotics commonly used for nuclear genetic engineering in green algae. These included the bleomycin/phleomycin family antibiotic zeocin, which intercalates into and cleaves DNA, and hygromycin and paromomycin which inhibit protein synthesis by disrupting translation. Whilst growth on agar was strongly inhibited by low concentrations of zeocin, higher concentrations of hygromycin and paromomycin were required to fully suppress growth (Supplementary Table 2). The *Streptoalloteichus hindustanus ble* gene (*Shble*) was identified as a potentially suitable selectable marker to progress *M. conductrix* SAG 241.80 genetic tool development. This gene confers resistance to bleomycin family antibiotics, including zeocin, and has been utilised for nuclear genetic engineering in other Trebouxiophyceae algae, including Chlorellaceae species (Yoshimitsu *et al*., 2018; Dahlin *et al*., 2019; Gonzalez-Esquer *et al*., 2019; Tatara *et al*., 2020; Gu *et al*., 2023; Kasai *et al*., 2024).

Two vectors were designed and synthesised with each containing a codon-optimised *Shble* gene flanked by *M. conductrix* SAG 241.80 endogenous *lhcII* regulatory sequences. In one vector the *Shble* gene contained no introns, whilst in the other the coding region was interrupted by repetitive insertions of the first intron of the *M. conductrix* SAG 241.80 *lhcII* gene. Colonies emerged under zeocin selection on all agar plates spread with populations electroporated with *Shble* vectors (Figure 2A). However, the number of colonies obtained per population was significantly higher when introns were present within the *Shble* gene (transformation efficiency: 157 colonies per µg of vector DNA; transformation frequency: 2.18 x 10^-5^ colonies per electroporated cell) than when the *Shble* coding sequence was uninterrupted by introns (transformation efficiency: 34 colonies per µg of vector DNA; transformation frequency: 4.67 x 10^-6^ colonies per electroporated cell) (z = 26.1, p < 0.001). As the *Shble* protein product has a 1:1 stoichiometric binding relationship with bleomycin family antibiotics (Gatignol *et al*., 1988), relative transformation efficiencies are correlated with transgene expression levels (Baier *et al*., 2020). Therefore, these results demonstrate that the first intron of the *lhcII* gene enhances transgene expression in *M. conductrix* SAG 241.80, at least when expression is under the control of *lhcII* flanking sequences. This led us to incorporate introns within all subsequent transgenes utilised in this study. Almost all transformant populations that were transferred to liquid media maintained the ability to grow in the presence of zeocin even after long-term culturing (approximately 10 months) in non-selective conditions (Figure 2B). This is consistent with stable integration of transgenes within the nuclear genome and suggests the potential for heterologous protein expression to be sustained in the absence of a selection pressure when *M. conductrix* SAG 241.80 transformants are introduced into *P. bursaria* host cells. Presence of the *Shble* transgene was verified for all transformants assessed by PCR (Figure 2C; the unedited gel image is displayed in Supplementary Figure 3).

**Figure 2:**
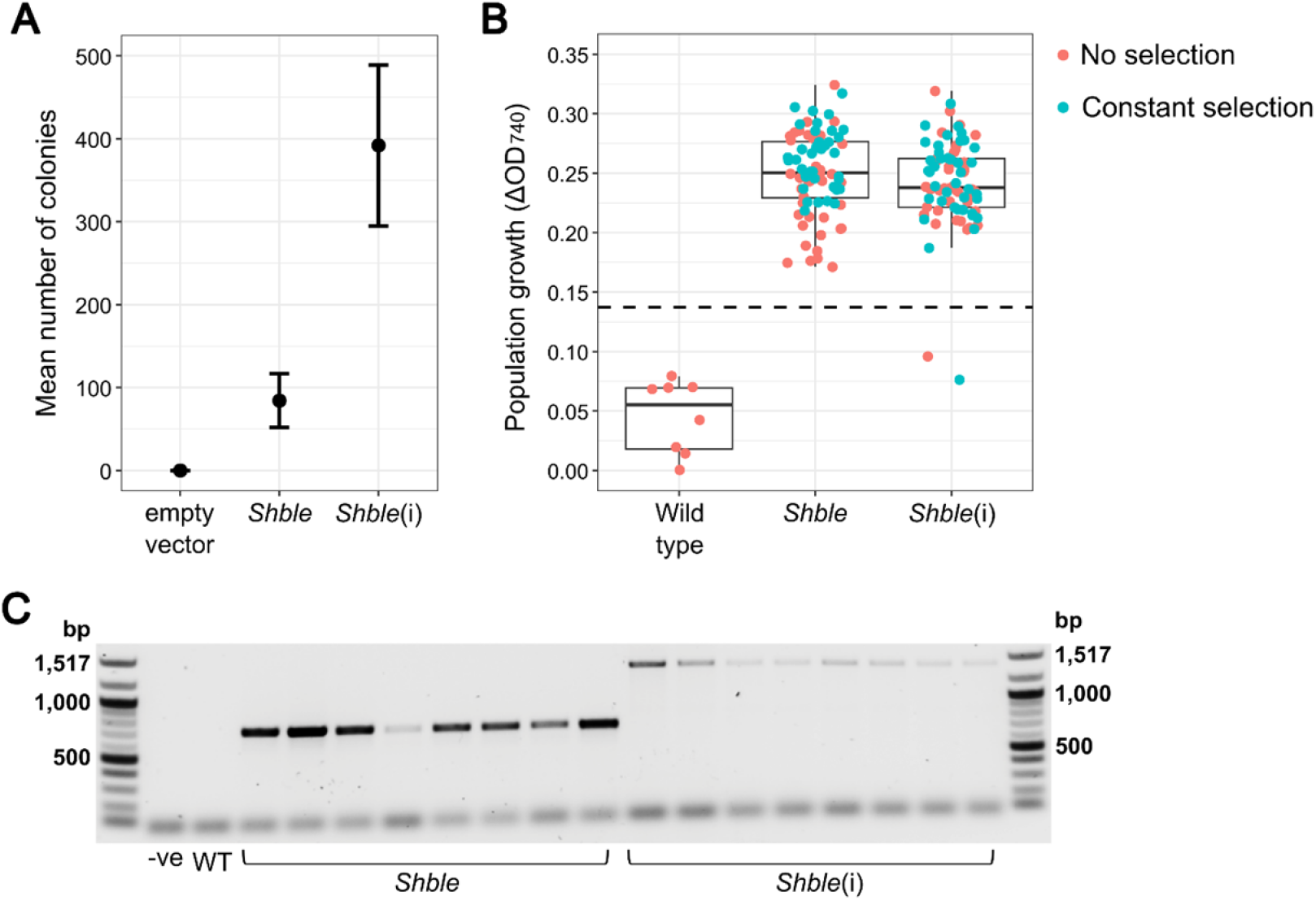
Expression of the *Shble* selectable marker gene is enhanced by endogenous introns. (A) Presence of endogenous introns within the *Shble* transgene (*Shble*(i)) enabled higher numbers of transformants to be isolated on selective agar plates, consistent with increased expression of the selection marker. Error bars represent standard errors of the means. (B) Population growth (ΔOD_740_, representing the change in optical density after 7 days) in selective media containing 5 µg/ml zeocin for *Shble* transformants which had been cultured with or without zeocin selection for approximately 10 months and wild type *M. conductrix* SAG 241.80 populations with no previous zeocin exposure (all wild type control populations died out during subculturing under zeocin conditions). Points represent replicate populations for the wild type strain and genotypes derived from distinct colonies for *Shble* transformants. The dashed line represents a threshold value of population growth (three standard deviations above the mean ΔOD_740_ observed for wild type controls) used to define zeocin resistance. Almost all transformants demonstrated zeocin resistance regardless of whether they had been maintained under constant selection or in the absence of a selection pressure for approximately 10 months, consistent with stable nuclear transgene integration. (C) Presence of the *Shble* transgene (and partial *lhcii* flanking sequences) was verified for all *Shble* and *Shble*(i) transformants assessed by PCR (-ve: no DNA template; WT: wild type *M. conductrix* SAG 241.80 genomic DNA template).

### An extended viral 2A peptide enables co-expression of multiple transgenes

Viral 2A peptides disrupt peptide bond formation during translation and result in production of two or more independent proteins from a single, polycistronic mRNA transcript. These have been used to couple expression of non-selectable genes to selectable markers in a range of green algae, including Trebouxiophyceae species, mitigating effects of transgene silencing and enabling transformant cell lines with desirable characteristics to be isolated without the need for extensive phenotype screening (Rasala *et al*., 2012, 2013; Plucinak *et al*., 2015; Dahlin *et al*., 2019; Gu *et al*., 2023). We assessed the capacity of an extended Foot-and-Mouth Disease Virus (FMDV) 2A peptide (Plucinak *et al*., 2015) to enable co-expression of *Shble* and a codon-optimised *mRuby3* fluorescent reporter gene in *M. conductrix* SAG 241.80. Vectors were designed and synthesised with transgenes placed in both possible orientations, i.e. one with *Shble* located upstream of the 2A peptide encoding sequence and *mRuby3* (*Shble*-2A-*mRuby3*) and one in the alternative orientation (*mRuby3*-2A-*Shble*). In both cases, the single open reading frames were flanked by *M. conductrix* SAG 241.80 endogenous *lhcII* regulatory sequences. We selected for transformants across a range of zeocin concentrations, predicting that higher concentrations could enable isolation of transgenic cell lines with stronger *mRuby3* expression because higher levels of *Shble* expression would be required for survival.

Fluorescent transgene expression was assessed by flow cytometry for a set of transformants obtained for each *mRuby3* vector and zeocin concentration by comparing mean chlorophyll normalised *mRuby3* fluorescence with controls carrying the *Shble* transgene only. No clear patterns associated with the concentration of zeocin imposed during colony isolation were observed (Figure 3A). However, whilst *mRuby3* expression was detectable in transformants obtained using both vectors, markedly superior fluorescent signals were observed when *mRuby3* was placed in the upstream position (Figure 3A). Over 90% of screened *mRuby3*-2A-*Shble* transformant populations displayed *mRuby3* expression levels above a detection threshold, with 84% showing at least a 100-fold increase in mean chlorophyll normalised *mRuby3* fluorescence relative to background levels in the controls. In contrast, *mRuby3* expression was detectable in < 70% of screened *Shble*-2A-*mRuby* transformant populations, and only ∼ 5% displayed at least a 100-fold increase in mean chlorophyll normalised *mRuby3* fluorescence relative to the controls. This demonstrates that gene orientation within 2A peptide-based constructs is an important determinant of transgene expression in *M. conductrix* SAG 241.80. Cytoplasmic localisation of *mRuby3* was clearly visible in a top *mRuby3*-2A-*Shble* transformant expression line by confocal fluorescence microscopy (Figure 3B). This is consistent with effective processing of the 2A peptide during translation and production of two independent proteins since the *Shble* protein product is predominantly confined to the nucleus in green algae (Fuhrmann *et al*., 1999; Plucinak *et al*., 2015).

**Figure 3:**
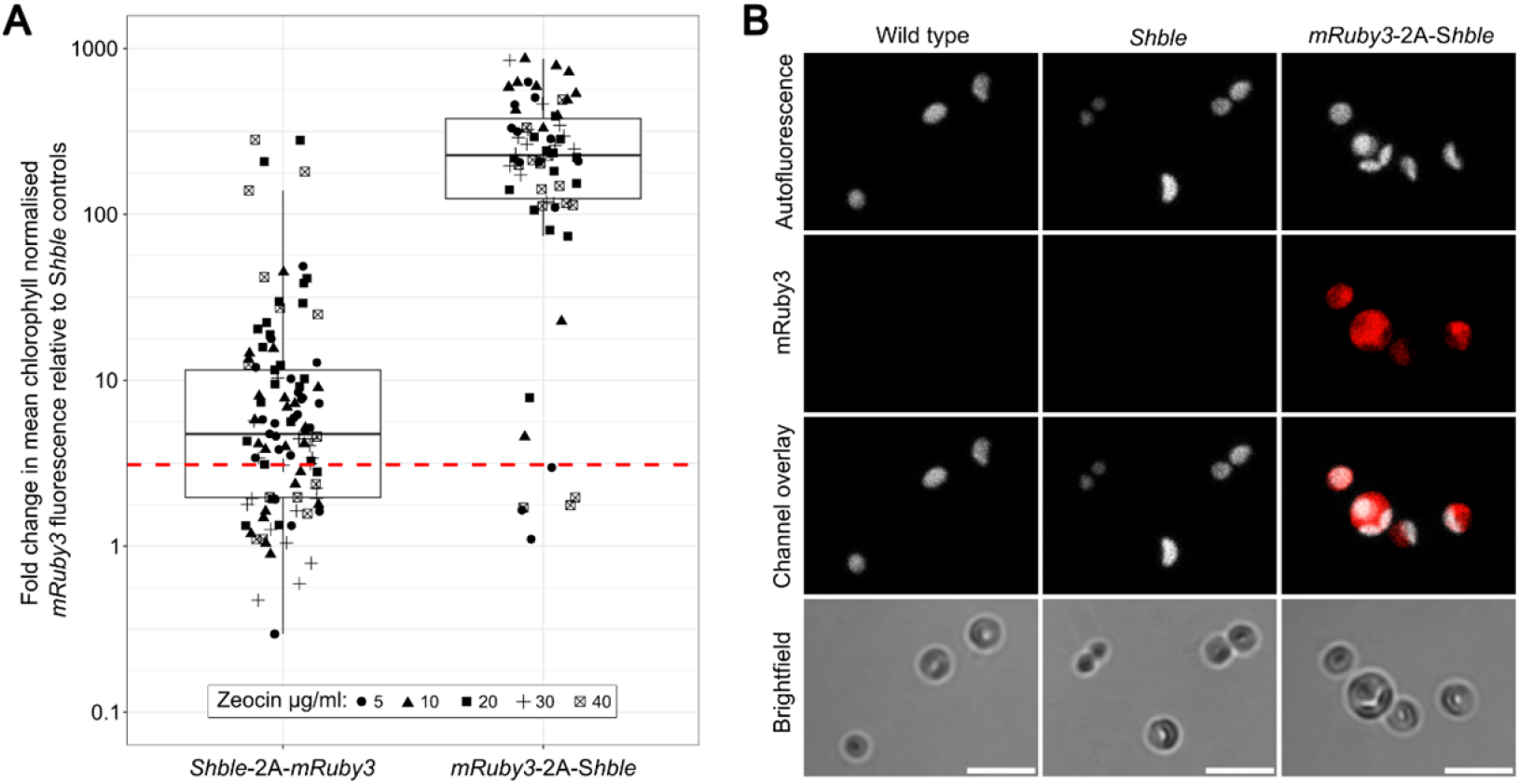
Transgene orientation within synthetic viral 2A peptide constructs is an important determinant of heterologous protein expression in *M. conductrix* SAG 241.80. (A) Fold change in mean chlorophyll normalised *mRuby3* fluorescence (based on flow cytometry data recorded for 10,000 cells per population) relative to controls (transformants carrying a *Shble* resistance cassette) for transformant genotypes obtained following electroporation with *Shble*-2A-*mRuby* or *mRuby3*-2A-*Shble* vectors and isolation at different zeocin concentrations. The dashed line represents the detection threshold for *mRuby3* expression (three standard deviations above the mean chlorophyll normalised *mRuby3* background signal for controls). Orientation within viral 2A peptide constructs clearly impacts the expression of non-selectable transgenes, with higher levels of expression occurring for transgenes placed upstream of a selectable marker. (B) Representative confocal microscopy images of *M. conductrix* SAG 241.80 wild type and transformant strains, demonstrating cytoplasmic localisation of *mRuby3* fluorescence in a top *mRuby3*-2A-*Shble* transformant expression line. Images were taken with a Zeiss LSM 980 confocal microscope using a 40X objective. Scale bars = 10 µm.

### Fluorescent transgene expression enables identification of distinct endosymbiont genotypes within host cells

We next introduced the same top *mRuby3*-2A-*Shble* transformant expression line into *P. bursaria* cells cleared of their native endosymbionts to determine if *mRuby3* expression could be detected in an endosymbiotic context in the absence of zeocin selection. Wild type *M. conductrix* SAG 241.80 cells and a zeocin-resistant transformant cell line carrying the *Shble* transgene were also introduced into cleared *P. bursaria* cells as controls. Expression of *mRuby3* was clearly detectable by confocal fluorescence microscopy three weeks after the *mRuby3*-2A-*Shble* transformant genotype was introduced into host cells (Figure 4A). Analyses of chlorophyll and mRuby3 channel images for *P. bursaria* cells containing this transformant genotype revealed a detectable *mRuby3* signal within almost every algal cell (Supplementary Figure 4A).

**Figure 4.**
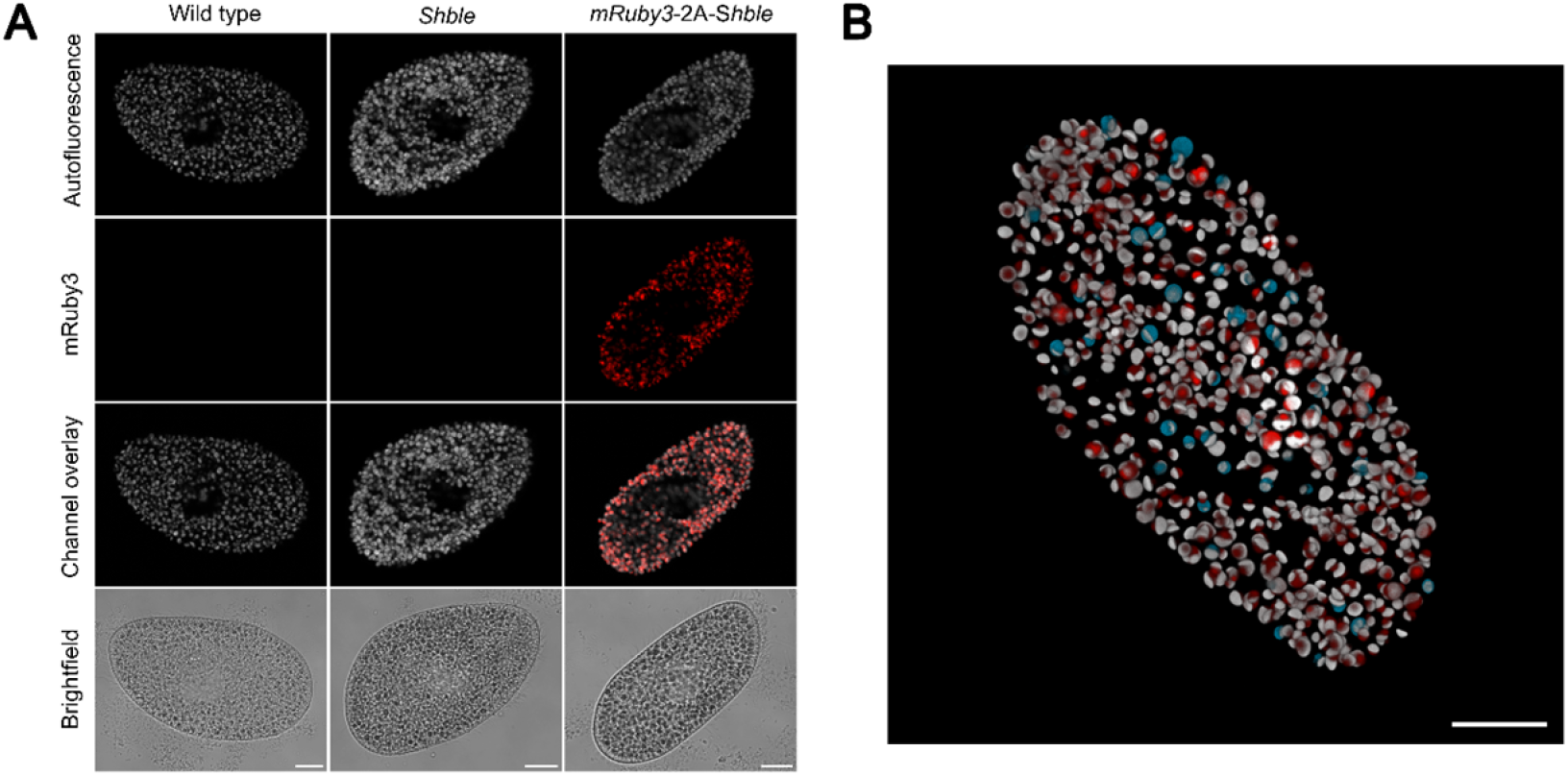
Fluorescent transgene expression enables identification of distinct genotypes within host cells. (A) Representative confocal microscopy images of *P. bursaria* host cells three weeks after introduction of the *M. conductrix* SAG 241.80 wild type, a zeocin resistant *Shble* transformant and a top *mRuby3*-2A-*Shble* transformant expression line. (B) 3D reconstruction (47 x Z-stacks) of algae within a *P. bursaria* cell 1 week after introduction of *mCerulean3*-2A-*Shble* and *mRuby3*-2A-*Shble* transformants. Chlorophyll autofluorescence is displayed in grey whilst blue and red represent *mCerulean3* and *mRuby3* fluorescence, respectively. Images were taken with a Zeiss LSM 980 confocal microscope using a 20X objective. Scale bars = 20 µm.

The homogeneity of *mRuby3* expression across *M. conductrix* SAG 241.80 cells suggests that it should be feasible to identify and quantify distinct transformant genotypes, specifically alternatively engineered variants, within an endosymbiotic context if they express different fluorescent proteins with spectral properties that can be distinguished by fluorescence microscopy. To test this, we transformed *M. conductrix* SAG 241.80 cells using a vector containing a codon-optimised *mCerulean3* transgene (*mCerulean3*-2A-*Shble*) and identified a strong expression line based on visual assessment by widefield fluorescence microscopy. This was co-introduced with a strong *mRuby3* expression line into *P. bursaria* cells cleared of their native endosymbionts. Z-stack images were collected 1 week after the introductions, then images were merged to generate 3D reconstructions for further analysis. The two distinct *M. conductrix* SAG 241.80 transformant genotypes could be unambiguously distinguished within host cells (Figure 4B), with 14.9 % of endosymbiont cells being assigned to the *mCerulean3* expression line and 79.2 % to the *mRuby3* expression line in the representative *P. bursaria* cell displayed (no clear assignment was possible for the remaining 5.9% of algal cells; Supplementary Figure 4B).

## Discussion

Our motivation for developing genetic tools for *M. conductrix* SAG 241.80 was to enable new approaches for addressing unsolved questions in photosymbiosis using the experimentally tractable *P. bursaria* model system. For applications to be useful in an endosymbiotic context, expression of transgenes must be stable in the absence of a selection pressure (assuming no natural resistance phenotype for the host) and sufficiently robust to enable observations within host cells. We have developed an efficient and reliable, stable transformation protocol for *M. conductrix* SAG 241.80 and identified properties of synthetic gene constructs which enable robust transgene expression. We found that transformed cells can be maintained for at least 10 months without loss of antibiotic resistance, even when cultured in non-selective conditions, and that coupling the expression of codon-optimised, intron-containing fluorescent transgenes to a selectable marker using a viral 2A peptide system enables isolation of transformants with sufficiently high levels of fluorescent protein expression for detection within the host environment. The principles established in this study provide a basis for further genetic tool development for *M. conductrix* SAG 241.80, for instance through construction of modular cloning resources and optimisation of targeted knock-out and/or knock-down approaches.

A key breakthrough during protocol development was the discovery that prolonged culturing in hyperosmotic conditions prior to electroporation dramatically increased population level *nanoluc* expression, an indicator of overall transformation success. This suggests that structural changes in the plasma membrane and/or cell wall during acclimation to hyperosmotic conditions may increase permeability to exogenous DNA and/or enhance post-electroporation viability in *M. conductrix* SAG 241.80, and reflects similar findings in gram-positive bacteria (Xue *et al*., 1999), fission yeast (Suga *et al*., 2003) and other Trebouxiophyceae algae (Tatara *et al*., 2020). We anticipate that using hyperosmotic growth conditions to prepare electrocompetent cells could facilitate genetic modification of Chlorellaceae species more generally, with implications for the development of heterologous expression systems for high value products (Yang *et al*., 2016; Chen and Ward, 2024).

We identified two synthetic construct design variables that had a large impact on transgene expression in *M. conductrix* SAG 241.80, including the presence or absence of introns within transgenes and orientation of transgenes in viral 2A peptide-based polycistronic sequences. It is well established that incorporation of introns can improve nuclear transgene expression in a range of eukaryotes, including *Chlamydomonas reinhardtii* (Lumbreras *et al*., 1998; Baier *et al*., 2018, 2020). Though mechanisms underpinning intron-mediated enhancement of gene expression remain unclear, this could be linked to the presence of intrinsic enhancer elements in specific intron sequences and/or positive feedback between spliceosome activity and transcription (Lumbreras *et al*., 1998; Rose and Beliakoff, 2000; Gallegos and Rose, 2015; Baier *et al*., 2018, 2020; Schroda, 2019). It has also been demonstrated that the presence of introns can suppress post-transcriptional transgene silencing in plants (Christie *et al*., 2011; Dadami *et al*., 2013), raising the possibility that a similar phenomenon could occur in green algae. *M. conductrix* genes are intron rich, with > 98% of genes containing introns and mean numbers of introns per gene ranging from 9.7 to 13.9 depending on the strain and gene prediction pipeline applied (Leonard *et al*., 2025b). Primary transcripts thus undergo intensive processing in *M. conductrix* SAG 241.80 and customising transgenes by insertion of introns at least partially mimics the architecture of endogenous genes.

Non-selectable genes are typically placed downstream of a selectable marker in 2A peptide-based constructs used for transgene expression in green algae (Rasala *et al*., 2012, 2013; Dahlin *et al*., 2019; Suttangkakul *et al*., 2019; Gu *et al*., 2023). However, we obtained a higher proportion of *mRuby3* positive transformants and observed considerably higher levels of fluorescence when *mRuby3* was placed in the upstream position. This could reflect differential impacts of truncations and/or rearrangements of the integrated exogenous DNA resulting from endogenous nuclease activity (Plucinak *et al*., 2015; Molina-Márquez *et al*., 2020). Reduced expression of *mRuby3* in the downstream position could also occur due to ribosomal drop-off, whereby the ribosome detaches prematurely from the mRNA transcript during translation (Liu *et al*., 2017), and/or instability resulting from the N-terminal addition of a proline residue derived from the 2A peptide C-terminal (Reinhardt *et al*., 2020). During translation of 2A peptide-based polycistronic mRNAs, the 2A peptide amino acids (excluding the proline residue mentioned above) are incorporated as C-terminal extensions to proteins encoded by the upstream gene (Donnelly *et al*., 2001). Many proteins will be robust to such C-terminal modifications, but in some cases this could lead to disrupted function, for instance when active sites are present within C-terminal domains (Plucinak *et al*., 2015). Although improved outcomes may not be a general rule, our results highlight the potential benefit that can be derived from placing genes of interest in the upstream position within 2A peptide-based constructs, particularly when genetic engineering endeavours are hampered by poor transgene expression.

The results described in this study provide foundations for generating *M. conductrix* SAG 241.80 transformants that are specifically tailored to explore mechanisms involved in the establishment and maintenance of photosymbiosis. For instance, using 2A peptide-based vectors as a platform to overexpress endogenous genes (Plucinak *et al*., 2015; Tokunaga *et al*., 2019), the functions of putative symbiosis associated genes could be assessed by examining phenotypic consequences of overexpression under ecologically relevant conditions. Moreover, as distinct *M. conductrix* SAG 241.80 genotypes can be identified based on fluorescent transgene expression, the colonisation abilities and fates of overexpression cell lines could be evaluated in direct relation to cells with endogenous expression levels by monitoring competition dynamics within the host environment. Such experiments could be conducted under different host RNAi treatments, allowing the endosymbiont and host metabolic interaction networks to be investigated simultaneously. Collectively, the methods developed here and elsewhere (He *et al*., 2019; Jenkins *et al*., 2021a, 2021b), combined with emerging genome data (Leonard *et al*., 2025a, 2025b), provide the tools to exploit *P. bursaria* as a system for exploring molecular and cellular functions that underpin endosymbiotic partner recognition, metabolic interactions, policing of symbiotic behaviours, and how the host selects variant algal genotypes for long-term partnership or destruction.

## Methods

### Culturing conditions

*M. conductrix* SAG 241.80 was obtained from the Culture Collection of Algae at Göttingen University (SAG), Germany. For standard culturing, *M. conductrix* SAG 241.80 was maintained in liquid modified Bold’s Basal Medium (MBBM: www.ccap.ac.uk) or on MBBM agar plates containing 1.5% bacteriological agar and 0.5% glucose, with a 14:10 L/D cycle at 24°C. *P. bursaria* 186b (CCAP 1660/18) cells were cultured in New Cereal Leaf-Prescott Liquid medium (NCL) supplemented with β-sitosterol and bacterized with *Klebsiella pneumoniae* SMC as previously described (Jenkins *et al*., 2021a). *P. bursaria* populations were maintained with a 12:12 L/D cycle at 20°C until being colonised with *M. conductrix* SAG 241.80 (or transformants derived from this strain), after which they were maintained with a 14:10 L/D cycle at 24°C.

### Antibiotic resistance assays

Sensitivity of *M. conductrix* SAG 241.80 to zeocin, hygromycin and paromomycin was determined by spotting serial dilutions of cells on to MBBM agar plates and categorising growth relative to no antibiotic controls after 4 weeks. Four replicate spots were monitored per antibiotic concentration and cell dilution. Initially, concentrations of 0, 5, 10, 20, 50 and 100 µg/ml were tested for each antibiotic. Sensitivity assays were then repeated for zeocin across a finer scale gradient of 0, 1, 2, 3, 4, and 5 µg/ml.

### Vector construction

Four *nanoluc* vectors were generated to assess the utility of different noncoding endogenous sequences for driving transgene expression. As promoter and terminator sequences have not been mapped for *M. conductrix* SAG 241.80, a *nanoluc* gene was fused between sequences of 1 kb in length which flank a set of nuclear encoded genes (ef1α, lhcii, PsaD, RBCS2, Supplementary Table 1). These were selected based on high expression levels (Arriola *et al*., 2018; NCBI accession number GSE98781) and/or utility of equivalent sequences in other green algal transformation studies. A codon usage table (Supplementary Table 3) was generated for highly expressed *M. conductrix* SAG 241.80 nuclear genes using the cusp tool in Emboss (Rice *et al*., 2000). As the pattern of codon usage was highly similar to the nuclear codon usage of *C. reinhardtii* (Kazusa codon usage database: www.kazusa.or.jp), we used a *nanoluc* gene sequence with demonstrated functionality which had been codon optimised for expression in *C. reinhardtii* (Crozet *et al*., 2018). Sequences were synthesised and inserted into pUC19 by Synbio Technologies using HindIII and EcoRI (ef1α and RBCS2) or HindIII and XbaI (lhcii and PsaD) cloning sites.

Two vectors were generated with a codon optimised *Shble* selectable marker gene fused between endogenous *M. conductrix* SAG 241.80 *lhcii* flanking sequences of 1 kb in length. In one version, the coding region contained repeated insertions of the first intron of the *M. conductrix* SAG 241.80 *lhcii* gene at positions selected using the Intronserter web tool (Jaeger *et al*., 2019). Three polycistronic vectors were then generated to couple fluorescent protein expression to *Shble* expression via an extended FMDV 2A peptide (Plucinak *et al*., 2015). Two contained codon-optimised *mRuby3* and *Shble* genes placed in alternative orientations, i.e. one with *mRuby3* upstream of the extended FMDV 2A peptide encoding sequence and *Shble* and the other vice versa. The third contained a codon optimised *mCerulean3* gene upstream of the extended FMDV 2A peptide encoding sequence and *Shble* gene. All transgenes in polycistronic vectors contained repeated insertions of the first intron of the *M. conductrix* SAG 241.80 *lhcii* gene at positions selected using the Intronserter web tool. Sequences were synthesised and inserted into a pUC19 plasmid backbone by Genscript using HindIII and either BamHI or XbaI cloning sites. Sequences of all DNA fragments utilized in this study are provided in Supplementary Table 1.

### Transformation

Here we describe an optimised transformation protocol which was developed by selecting conditions that maximised population level *nanoluc* expression (see results section). Vectors were linearised by digestion with HindIII-HF (New England Biolabs), then purified using a Wizard® SV Gel and PCR Clean-Up System (Promega) and eluted in nuclease free water. *M. conductrix* SAG 241.80 populations were harvested in mid log phase after 48 hrs growth in MBBM containing 0.1 M MOPS (pH 7 ± 0.1) by centrifugation at 2,800 x g for 5 mins. Cell densities were determined using a Countess II FL Automated Cell Counter (Invitrogen) fitted with a Cy5 EVOS^TM^ light cube. Cells were pelleted by centrifugation as above, washed in 1 ml ice-cold MAX Efficiency^TM^ Transformation Reagent for Algae (Invitrogen), then resuspended in ice-cold MAX Efficiency^TM^ Transformation Reagent for Algae supplemented with 40 mM sucrose at a density of approximately 4 x 10^8^ cells/ml. Cell suspensions were kept on ice then, for each transformation, 45 µl of cell culture was mixed with 5 µl of linear vector (0.5 µg/µl) and transferred to an ice-cold 2 mm electroporation cuvette (VWR). Cells were electroporated with a 5.0 msec, 8.5 kV/cm square-wave pulse using a Gene Pulser Xcell^TM^ electroporation system (Bio-Rad), then were incubated on ice for 2-4 mins. Electroporated cells were transferred into 24 well plates in a final volume of 2 ml MBBM and allowed to recover during an extended dark period of 16 hrs. For *nanoluc* transformations, cells were incubated with a 14:10 L/D cycle for an additional 48 hrs before being processed as described below. For all other transformations, cells were incubated in the light for approximately 6 hrs then were plated on selective agar plates containing 5 µg/ml zeocin (except where stated otherwise) and supplemented with 100 µg/ml ampicillin to prevent occurrence of bacterial contamination.

### Nanoluc assays

Nanoluc luciferase assays were performed on cell lysates to quantify population level transgene expression using a Nano-Glo® Luciferase Assay System kit (Promega). Populations were pelleted by centrifugation at 2,800 x g for 5 mins, resuspended in 50 µl Nano-Glo® Luciferase Assay Buffer with a small quantity of acid washed 212-300 µm glass beads (Sigma-Aldrich), and homogenised using a TissueLyser II (Qiagen) for 5 minutes at the maximum speed (30 Hz). Cell debris was pellet by centrifugation at 14,000 x g for 10 mins then supernatants were transferred to a Greiner CELLSTAR® white 96 well plate with an equal volume of Nano-Glo® Luciferase Assay Reagent. Luminescence was measured using a Varioskan LUX multimode microplate reader (Thermo Scientific).

Effect sizes were calculated for each treatment as natural log-transformed response ratios (lnRRs) relative to background luminescence detected for controls electroporated without vector DNA. For a given treatment, lnRR = ln(*X*_T_/*X*_C_), where *X*_T_ and *X*_C_ represent mean luminescence values recorded for treatment populations (n ≥ 4) and control populations (n ≥ 4), respectively. Standard errors were calculated as previously described (Hedges *et al*., 1999). Treatment effect sizes were displayed as fold change values relative to controls to simplify interpretation and were considered as significantly different from controls if 95% confidence intervals did not overlap with one.

When comparing transformation conditions during protocol development (see results section), we occasionally observed high variability in population density prior to Nanoluc luciferase assays, suggesting differences in post-electroporation viability and/or growth rates across treatments. However, neither cell densities nor protein concentrations were equalised before performing the assays because our objective was to identify conditions which maximised population level transgene expression regardless of whether this was achieved with high transformation rates but low viability, low transformation rates with high viability, or somewhere in between.

### Characterising *Shble* transformants

Selective agar plates were imaged using a Nikkon D7000 camera three weeks after plating with *M. conductrix* SAG 241.80 populations that had been electroporated with an empty pUC19 vector or a *Shble* vector with or without introns. Colonies were counted with ImageJ (Schneider *et al*., 2012) using the Analyse Particles tool and counts were compared using a generalised linear model with a poisson error distribution in R version 4.4.3.

Zeocin resistant colonies obtained for each *Shble* vector and wild type colonies from a standard MBBM plate (since no colonies emerged under zeocin selection for empty vector control populations) were transferred to a 96 well plate containing 200 µl MBBM per well. This stock plate was used to initiate populations in standard MBBM and in MBBM supplemented with 5 µg/ml zeocin, allowing growth to be monitored in each condition. Populations were maintained by transferring 10 µl of culture into fresh media approximately every 2 weeks. Transgene stability was assessed after approximately 10 months by monitoring growth in MBBM supplemented with 5 µg/ml zeocin based on optical density measurements at 740 nm (OD_740_) recorded using a FLUOstar Omega (BMG Labtech) plate reader.

Presence of the *Shble* expression cassette was assessed by PCR using primers which bind within *lhcii* regulatory sequences (primer sequences are provided in Supplementary Table 1). To extract genomic DNA, 50 µl of cell culture was pelleted by centrifugation at 2,800 x g for 5 mins then pellets were resuspended in 50 µl nuclease free water with a small quantity of acid washed 212-300 µm glass beads (Sigma-Aldrich). The cells were incubated at 95°C for 5 mins, vortexed, and then centrifuged at 14,000 x g for 5 mins. The supernatant was used as DNA template in PCRs that were performed using GoTaq® Green Master Mix (Promega) supplemented with 4 mM MgCl_2_ and 5% DMSO following the manufacturer’s recommended protocol with an annealing temperature of 58°C and an extension time of 1 min 20 secs. This extension time was not sufficient for amplification of the endogenous genomic region containing the *lhcii* gene and flanking sequences (primer binding sites flank an approximately 2.7 kb region).

### Detection of fluorescent transgene expression

*M. conductrix* SAG 241.80 transformants carrying *Shble*-2A-*mRuby3* or *mRuby3*-2A-*Shble* expression cassettes were examined by flow cytometry and compared with zeocin resistant controls transformed with an *Shble* only vector. Zeocin resistant colonies were inoculated into liquid MBBM supplemented with 5 µg/ml zeocin and, after one subculture, log phase populations were examined for evidence of *mRuby3* expression. Data were recorded for 10,000 cells per population using a cytoFLEX LX flow cytometer (Beckman Coulter), with *mRuby3* expression detected using the 561 nm laser and a 585/42 nm bandpass filter and chlorophyll autofluorescence detected using the 488 nm laser and a 690/50 nm bandpass filter. Data were analysed with FlowJo v10 (BD Life Sciences) using a nested gating strategy to identify single, chlorophyll positive cells. *mRuby3* fluorescence was divided by chlorophyll fluorescence to obtain chlorophyll normalised values for each cell then mean chlorophyll normalised *mRuby3* fluorescence values for each transformant population were used to determine fold change relative to background levels observed for *Shble* controls. We defined the detection threshold as three standard deviations above the mean background signal observed for *Shble* controls.

*M. conductrix* SAG 241.80 transformants carrying *mCerulean3*-2A-*Shble* expression cassettes were visually assessed for levels of *mCerulean3* fluorescence and compared with controls transformed with an *Shble* only vector using an Olympus CKX53 inverted widefield microscope with a blue 475/30 nm excitation filter.

### Host introductions

*P. bursaria* 186b cells were cleared of their native *M. conductrix* 186b algal endosymbionts (∼ 99.9% nucleotide identity to *M. conductrix* SAG 241.80; Leonard *et al*., 2025b) by treatment with 10 µg/ml paraquat dichloride hydrate for eight days followed by 10 µg/ml cycloheximide for six days. Clearance was confirmed by visual assessment using an Olympus CKX53 inverted widefield microscope. Cleared host cells were collected in pluriStrainer cell strainers (pluriSelect) with a 20 µm pore size and washed with NCL to remove traces of the treatments. Log phase populations of wild type and transformant *M. conductrix* SAG 241.80 genotypes were concentrated by centrifugation at 2,800 x g for 5 mins, washed then resuspended in NCL and incubated with cleared host cells for 24 hours at a ratio of approximately 1:100,000 host:algal cells. Host cells with introduced algae were collected in pluriStrainer cell strainers as above and washed with NCL to remove the majority of aposymbiotic algae. They were then resuspended in fresh NCL bacterized with *K. pneumoniae* and incubated for at least one week before imaging. Cleared *P. bursaria* 186b host cells with no algae introduced were monitored throughout this period as controls to ensure that no recolonisation of native algae occurred.

### Confocal laser scanning microscopy

Cells were imaged with an LSM 980 microscope (Zeiss) using a 40X objective for aposymbiotic algae and a 20X objective for endosymbiotic algae within host cells. Chlorophyll autofluorescence was excited with a 639 nm laser (aposymbiotic algae: 30% power and 850 V gain; endosymbiotic algae: 1.5% power and 650 V gain) and emission was detected between 649-694 nm. *mRuby3* was excited with a 561 nm laser (aposymbiotic algae: 40% power and 950 V gain; endosymbiotic algae: 2% power and 650 V gain) and emission was detected between 569-631 nm. *mCerulean3* was excited with a 445 nm laser (2.5% power and 750 V gain) and emission was detected between 450-550 nm. Images were processed and analysed using ImageJ (Schneider *et al*., 2012). Background fluorescence was removed from each fluorescence channel using the inbuilt ‘Subtract Background’ function and, where applicable, a 2D image was created by summarizing all z-stacks. To quantify the number of algae cells for each 2D fluorescence channel image, a nuclei segmentation was performed using the ‘StarDist’ plugin (Schmidt *et al*., 2018). Different thresholding algorithms were then tested on the segmented cell map and the algorithm with the best coverage was selected to create a counting mask (Yen *et al*., 1995).

## Acknowledgements

We thank David Booth at the Department of Biochemistry and Biophysics, University of California, San Francisco, for helpful suggestions regarding protocol development. We also gratefully acknowledge Robert Hedley and Vasiliki Tsioligka at the Don Mason Facility of Flow Cytometry, Sir William Dunn School of Pathology, University of Oxford, and the Micron Advanced Bioimaging Facility, University of Oxford (supported by Wellcome Strategic Awards 091911/B/10/Z and 107457/Z/15/Z) for their support and assistance in this work. This work was supported by the following grants from the Gordon and Betty Moore Foundation: ‘New Tools for Advancing Model Systems in Aquatic Symbiosis’ (GBMF9353) and ‘New Methods and Resources for Endosymbiosis and Eukaryogenesis Research’ (GBMF11490), as well as by a Royal Society University Research Fellowship to TAR (UF130382).

