## Supplementary materials for "Stable transformation of *Micractinium conductrix* SAG 241.80: New tools for exploring photosymbiotic interactions"

### Supplementary Table 1: Vector insert sequences and primer sequences

### Supplementary Table 2: Sensitivity to hygromycin, paromomycin and zeocin.

| | | Concentration (µg/ml) | | | | | | | | |
|---|---|---|---|---|---|---|---|---|---|---|
| **Antibiotic** | Density | 1 | 2 | 3 | 4 | 5 | 10 | 20 | 50 | 100 |
| **Hygromycin** | High | ND | ND | ND | ND | +++ | +++ | +++ | +++ | + |
| | Medium | ND | ND | ND | ND | +++ | +++ | +++ | + | - |
| | Low | ND | ND | ND | ND | +++ | +++ | ++ | - | - |
| **Paromomycin** | High | ND | ND | ND | ND | +++ | +++ | +++ | - | - |
| | Medium | ND | ND | ND | ND | +++ | +++ | +++ | - | - |
| | Low | ND | ND | ND | ND | +++ | +++ | ++ | - | - |
| **Zeocin** | High | ++ | ++ | + | - | - | - | - | - | - |
| | Medium | + | + | - | - | - | - | - | - | - |
| | Low | + | - | - | - | - | - | - | - | - |

+++ Equivalent to controls, ++ moderate/strong reduction relative to controls, + few isolated colonies, - no growth, ND not determined.

Serial dilution densities (25 µl aliquots per spot): high ~ 1 x $10^7$ cells/ml, medium ~ 1 x $10^6$ cells/ml, low ~ 1 x $10^5$ cells/ml.

### Supplementary Table 3

| Codon | AA | Fraction | Frequency | Number | Codon | AA | Fraction | Frequency | Number |
|---|---|---|---|---|---|---|---|---|---|
| **GCA** | A | 0.033 | 3.635 | 34 | **CCA** | P | 0.013 | 0.641 | 6 |
| **GCC** | A | 0.552 | 59.974 | 561 | **CCC** | P | 0.771 | 38.807 | 363 |
| **GCG** | A | 0.272 | 29.506 | 276 | **CCG** | P | 0.117 | 5.88 | 55 |
| **GCT** | A | 0.143 | 15.501 | 145 | **CCT** | P | 0.1 | 5.025 | 47 |
| **TGC** | C | 1 | 11.118 | 104 | **CAA** | Q | 0.003 | 0.107 | 1 |
| **TGT** | C | 0 | 0 | 0 | **CAG** | Q | 0.997 | 33.034 | 309 |
| **GAC** | D | 0.91 | 37.845 | 354 | **AGA** | R | 0 | 0 | 0 |
| **GAT** | D | 0.09 | 3.742 | 35 | **AGG** | R | 0.015 | 0.962 | 9 |
| **GAA** | E | 0.017 | 0.855 | 8 | **CGA** | R | 0.002 | 0.107 | 1 |
| **GAG** | E | 0.983 | 49.711 | 465 | **CGC** | R | 0.793 | 50.78 | 475 |
| **TTC** | F | 0.789 | 31.537 | 295 | **CGG** | R | 0.125 | 8.018 | 75 |
| **TTT** | F | 0.211 | 8.446 | 79 | **CGT** | R | 0.065 | 4.169 | 39 |
| **GGA** | G | 0.019 | 1.71 | 16 | **AGC** | S | 0.326 | 17.212 | 161 |
| **GGC** | G | 0.892 | 80.607 | 754 | **AGT** | S | 0.006 | 0.321 | 3 |
| **GGG** | G | 0.027 | 2.459 | 23 | **TCA** | S | 0.012 | 0.641 | 6 |
| **GGT** | G | 0.062 | 5.559 | 52 | **TCC** | S | 0.411 | 21.702 | 203 |
| **CAC** | H | 0.906 | 16.464 | 154 | **TCG** | S | 0.107 | 5.666 | 53 |
| **CAT** | H | 0.094 | 1.71 | 16 | **TCT** | S | 0.138 | 7.27 | 68 |
| **ATA** | I | 0.003 | 0.107 | 1 | **ACA** | T | 0.004 | 0.214 | 2 |
| **ATC** | I | 0.826 | 31.965 | 299 | **ACC** | T | 0.842 | 40.411 | 378 |
| **ATT** | I | 0.171 | 6.628 | 62 | **ACG** | T | 0.085 | 4.062 | 38 |
| **AAA** | K | 0.012 | 0.962 | 9 | **ACT** | T | 0.069 | 3.314 | 31 |
| **AAG** | K | 0.988 | 82.318 | 770 | **GTA** | V | 0.003 | 0.214 | 2 |
| **CTA** | L | 0.001 | 0.107 | 1 | **GTC** | V | 0.263 | 20.633 | 193 |
| **CTC** | L | 0.161 | 13.791 | 129 | **GTG** | V | 0.685 | 53.774 | 503 |
| **CTG** | L | 0.802 | 68.848 | 644 | **GTT** | V | 0.049 | 3.849 | 36 |
| **CTT** | L | 0.025 | 2.138 | 20 | **TGG** | W | 1 | 12.08 | 113 |
| **TTA** | L | 0 | 0 | 0 | **TAC** | Y | 0.906 | 24.802 | 232 |
| **TTG** | L | 0.011 | 0.962 | 9 | **TAT** | Y | 0.094 | 2.566 | 24 |
| **ATG** | M | 1 | 22.878 | 214 | **TAA** | * | 0.5 | 2.673 | 25 |
| **AAC** | N | 0.977 | 36.455 | 341 | **TAG** | * | 0.04 | 0.214 | 2 |
| **AAT** | N | 0.023 | 0.855 | 8 | **TGA** | * | 0.46 | 2.459 | 23 |

Coding GC 65.78%, 1st letter GC 61.92%, 2nd letter GC 43.98%, 3rd letter GC 91.45%

Codon usage table for the 50 most highly expressed *M. conductrix* SAG 241.80 genes generated using the cusp tool in Emboss.

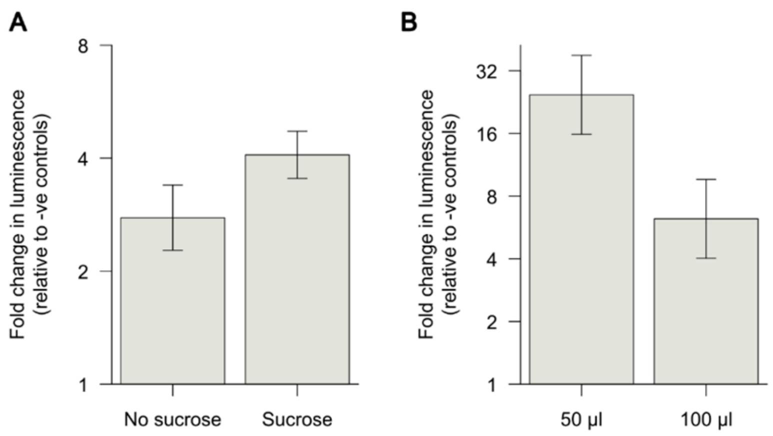


**Supplementary Figure 1: Development and refinement of a transformation protocol based on population level expression of NanoLuc™ luciferase.** (A) Supplementing the electroporation buffer with 40 mM sucrose and (B) reducing the volume of cell suspension used during electroporation whilst keeping total cell numbers constant improved population level expression of NanoLuc™ luciferase. Data are presented as fold change relative to controls electroporated without vector DNA and error bars represent 95% confidence intervals. Effects were considered as significantly different from controls when confidence interval error bars did not overlap with 1.

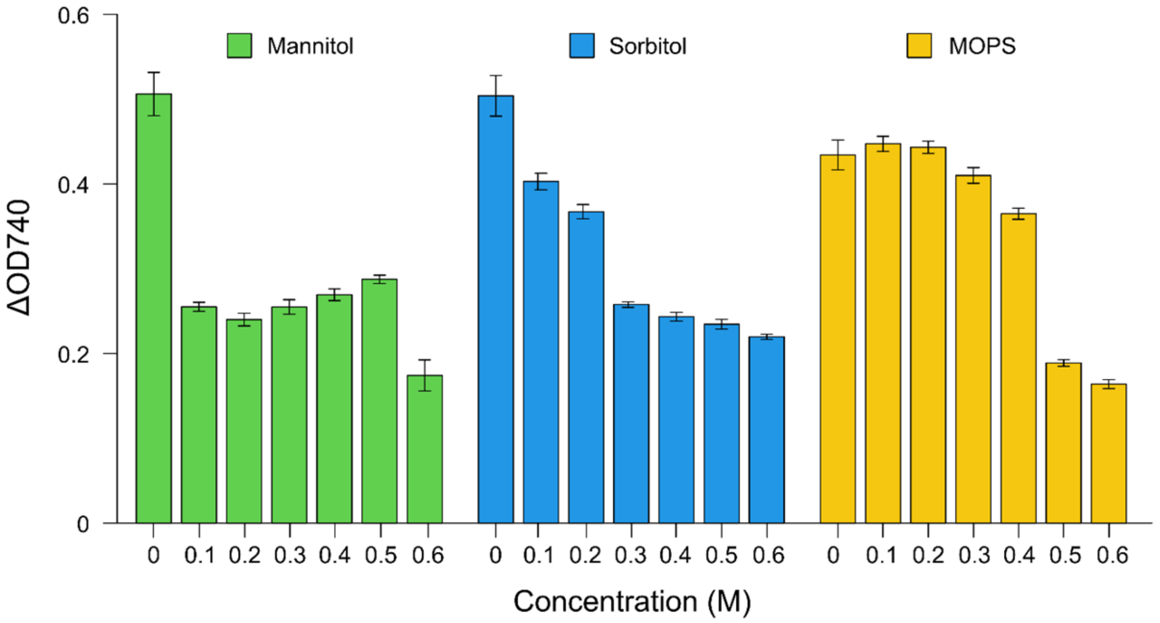


**Supplementary Figure 2: Sensitivity to mannitol, sorbitol and MOPS.** Population growth was determined based on changes in optical density ($\Delta OD_{740}$) after 7 days exposure to MBBM media containing different concentrations of mannitol, sorbitol or MOPS (3-(N-Morpholino)propanesulfonic acid).

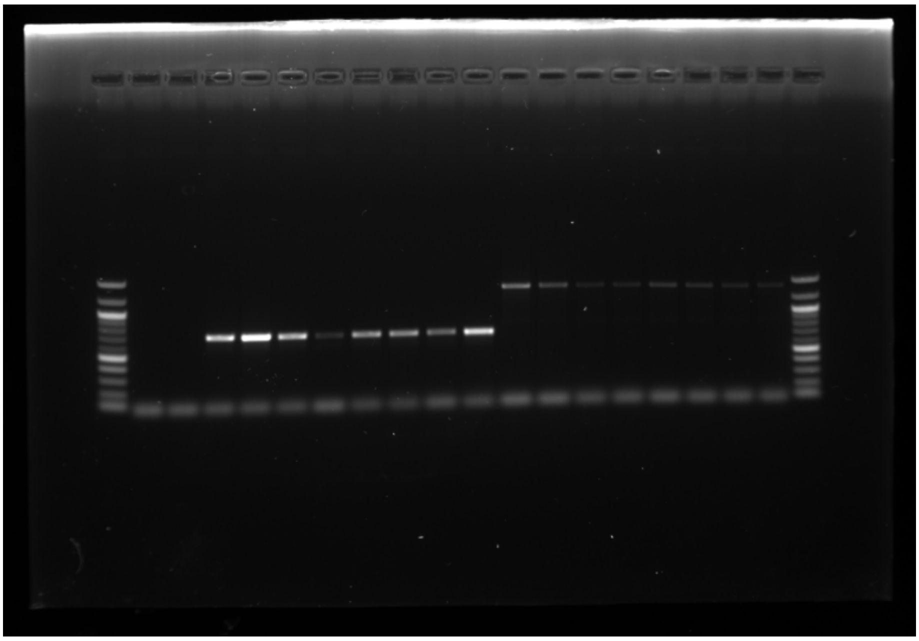

**Supplementary Figure 3: Unedited and uncropped gel image.** Bands verify the presence of the *Shble* transgene (+/- introns) for a selection of transformant cell lines. This image was used to generate Figure 3C.

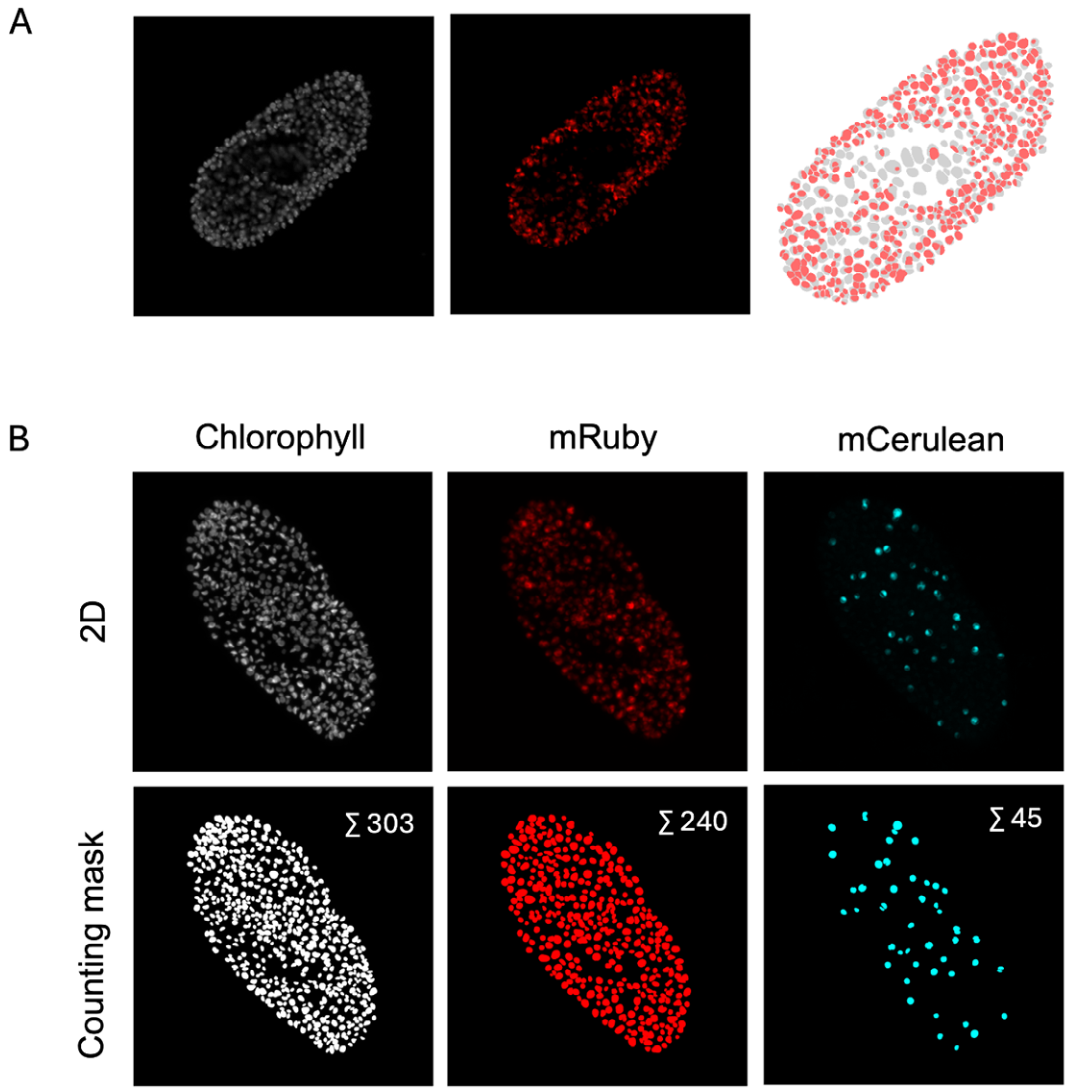


**Supplementary Figure 4: Confocal image analyses revealed homogeneous transgene expression within the host environment.** (A) An *mRuby3* signal was detectable in the majority of *mRuby3*-2A-*Shble* algae imaged within a *Paramecium bursaria* cell on a single plane (the proportion of algae with detectable *mRuby3* fluorescence is not reported due to deemed inaccuracies in the overlayed counting masks caused by detection of fluorescence from different planes). From left to right: chlorophyll autofluorescence (grey), *mRuby3* (red) and overlay of counting masks for each channel. (B) An *mCerulean3* or *mRuby3* signal was detectable in > 90% of transformant cells imaged within a *P. bursaria* cell based on multiple z-stack acquisition, demonstrating the capacity to assign endosymbiont identities in mixed-genotype interactions. The top row shows the summarized 2D image of all z-stacks of the different fluorescence channels. The bottom row shows the final counting masks used for quantification of the two transformation expression lines. Images were taken with a Zeiss LSM 980 confocal microscope using a 20X objective.
